# Identification of druggable targets from the interactome of the Androgen Receptor and Serum Response Factor pathways in prostate cancer

**DOI:** 10.1101/2024.08.15.608046

**Authors:** Haleema Azam, Colin Veale, Kim Zitzmann, Simone Marcone, William M. Gallagher, Maria Prencipe

## Abstract

**Objectives:** The Androgen Receptor (AR) is crucial for prostate cancer (PCa) progression. Despite the introduction of second-generation AR antagonist, majority of patients develop resistance. The Serum Response Factor (SRF) was identified as a player involved in a crosstalk with AR signalling pathway and associated resistance. Elevated SRF levels in PCa patients were associated with disease progression and resistance to enzalutamide. However, the molecular mediators of the crosstalk between SRF and AR still need to be elucidated. The objective of this study was to identify common interactors of the AR/SRF crosstalk as therapeutic targets.

**Methods:** Here we used affinity purification mass spectrometry (MS) to identify proteins that interact with both SRF and AR.

**Results:** Seven common interactors were identified, including HSP70, HSP0AA1, HSP90AB1, HSAP5, PRDX1 and GAPDH. Pathway analysis revealed that the PI3k/AKT pathway was the most enriched in the AR/SRF network. Moreover, pharmacological inhibition of several proteins in this network, including HSP70, HSP90, PI3k and AKT, significantly decreased cellular viability of PCa cells.

**Conclusions:** This study identified a list of AR/SRF common interactors that represent a pipeline of druggable targets for the treatment of PCa.

## Introduction

Castrate resistant prostate cancer (CRPC) grows independently of androgens. However, AR still plays a central role in driving progression to CRPC. Despite the addition of enzalutamide and other second-generation inhibitors of androgen biosynthesis, such as abiraterone acetate, for the treatment of advanced PCa, majority of patients develop resistance^1^. As new avenues of resistance to AR antagonists emerge, understanding AR’s relationship with co-regulators will aid in finding targets to disrupt the AR pathway in CRPC. The study of the crosstalk between AR and other signalling pathways involved in the plasticity of AR transcriptional network will shed light on the molecular mechanism behind the transition from androgen-sensitive to CRPC. One way in which AR co-factors control its transcriptional activity is through regulating AR nuclear translocation^2^. SRF is a transcription factor that plays a key role in cytoskeleton organisation and is implicated in proteins’ sub-cellular trafficking. Several studies have shown a crosstalk between SRF and AR^3–5^. In an isogenic model of CRPC, downregulation of SRF in the presence of DHT, stimulated an increase in AR transcriptional activity in the LNCaP Abl (androgen-independent), but not in the LNCaP parental cells (androgen-dependent), suggesting a negative feedback loop in the androgen-independent subline^4^. This negative feedback loop was also observed in patients’ tissues from CRPC bone metastases, where a negative correlation occurred between AR and SRF protein expression^4^. Another study supporting the AR/SRF relationship showed that AR and SRF shared a gene signature of 158 genes that were androgen responsive in LNCaP and VCaP cell lines^6^. This gene signature was associated with poor outcome in patients^6^. Other studies on the relationship between AR and SRF, showed that inhibition of protein kinase N1 (PKN1), an SRF co-factor responsible for androgen mediated SRF transcriptional activity, led to decreased expression of SRF transcriptional targets and increased expression of AR transcriptional targets^5^. Additionally, Four and Half Lim domain 2 (FHL2), a protein that regulates AR activity^7,8^, is under the transcriptional control of SRF^9^. In line with these data, we have demonstrated that elevated SRF expression is associated with enzalutamide resistance in patients^10^ and that inhibition of SRF reduces AR translocation to the nucleus^10^. Despite all the evidence supporting the crosstalk between SRF and AR, little is known about how SRF influences AR signalling and *vice versa*. Here we used co-immunoprecipitation (co-IP) and MS to identify common interactors of SRF and AR.

## Materials and Methods

### Cell culture and Reagents

LNCaP Parental cell line was routinely cultured in Advanced RPMI-1640 supplemented with 10% Fetal Bovine Serum (FBS), 100μl/mL streptomycin/100 U/mL penicillin and 1% Hepes. The isogenic LNCaP Abl subline was generated as previously described^11^ and routinely cultured like the parental LNCaP but replacing FBS with Charcoal Stripped Serum. Cell lines were maintained at 37°C in a humidified atmosphere and were routinely tested for mycoplasma. (5a,17b)-17-Hydroxy-androstan-3-one (DHT) was purchased from Sigma. JG-98, Ver-155008, Ganetespib, Ipatasertib and Alpesilib were purchased from MedChem Express.

### Small-interfering RNA (siRNA) and plasmids’ transfections

Two hundred thousand LNCaP Abl cells per well were seeded in 6 well plates and 1,500,000 cells in 10cm^3^ petri dishes. The following day, cells were transfected with siGENOME SMART pool targeting SRF, AR or siControl siRNA (all from Dharmacon), at a final concentration of 5nM, or with p-CGN SRF plasmid (Addgene plasmid 11977) or with p-HM6 empty vector at a final concentration of 1μg/μL, using lipofectamine 2000 (Invitrogen).

### Co-Immunoprecipitation Assay

Prior to cellular lysis, the antibody solution was prepared adding 2μg of either SRF (Novus, Biotechne), AR (Santa Cruz, California, US), Ms IgG or Rb IgG to 20μL A/G protein beads (Pierce) and 300μL PBS (Gibco). The beads/antibody mixture was incubated for an hour on a rotator at 4°C. Following incubation, the mixture was washed with ice-cold lysis buffer 3 times. Cells were scraped with 500μL of lysis buffer (500μL of 1% Triton x100, 1mL of 20mM Tris-HCl pH7.5, 1.5mL of 150mM NaCl and 50μL of 1mM MgCl_2_) and incubated for 10 mins on ice. 1mg of protein was added to the beads/antibody mix and incubated for 1 hour on a rotator at 4°C. Samples were washed 3 times in ice-cold lysis buffer.

### Peptide elution and digestion

Following Co-IP, the peptides in each sample were eluted with 60μL of ice-cold Elution Buffer I (0.012g Urea, 50μL of 1M Tris-HCl pH7.5) and 5μg/mL Trypsin (Promega, Seq Grade Modified) for 30 mins at RT. Samples were then centrifuged at 3000rpm for 30s. The supernatant was collected into a new Eppendorf tube, and 20μL of Elution Buffer II was added to each sample. This step was repeated twice. The supernatant was collected into a new centrifuge tube with a total volume of 110μL. Samples were left to digest overnight at 37°C at 300rpm.

### Liquid Mass spectrometry (Bruker timsTOF Pro) and data analysis with MaxQuant

A Bruker timsTOF Pro MS (Bruker Daltonics, Bremen, Germany) connected to an EvoSep One chromatography system was used. The timsTOF Pro MS was run using positive ion polarity with TIMS (Trapped Ion Mobility Spectrometry) and PASEF (Parallel Accumulation Serial Fragmentation) modes. Accumulating ramp times for the TIMS were set at 100ms, with the ion mobility ranging from 0.6 to 1.6 Vs/cm. A mass range from 100 to 1,700 m/z was the set range to record the spectra of ions. The precursor MS Intensity Threshold was set to 2,500 and the precursor Target Intensity was set to 20,000. Each PASEF cycle included one MS ramp for precursor detection, accompanied by 5 PASEF MS/MS ramps and a total cycle time of 1.03s. Peptides were separated using reverse-phase C_18_ Endurance column (15cm x 150µm ID, C18, 1.9µm) using the Evosep pre-set 30 SPD method. Mobile phase A consisted of 0.1% (v/v) formic acid in water and phase B included 0.1% (v/v) formic acid in acetonitrile. Peptides were separated by increasing gradient of solvent B for 44 minutes with a flow rate of 0.5µL/min. MaxQuant v1.6.17.0 was used by applying the *Homo sapiens* subset of the Uniprot Swissprot database against the raw data. A contaminants database was included in the search and the ‘Match Between Runs’ and ‘Label free quantification’ were selected^12^. The minimum peptide length allowed was 7 amino acids. False discovery rate (FDR) for peptides was set at 0.01. Protein intensity of each identified protein was normalised to obtain the label free quantification intensity (LFQi) value. A ProteinGroups.txt output file generated by MaxQuant was used for subsequent data analysis.

The mass spectrometry proteomics data have been deposited to the ProteomeXchange Consortium via the PRIDE partner repository with the dataset identifier PXD051524.

### Perseus analysis

The ProteinGroups.txt file generated by MaxQuant was processed on Perseus (v1.6.12.0)^13^. Data was filtered based on the LFQi values, and possible contaminants and reverse protein hits were removed from the list of identified proteins. Proteins that were not present in at least two samples were excluded from the analysis. The data was then imported onto Excel, where the LFQi ratio between the AR endogenous/siRNA or SRFvector/siRNA were calculated. Proteins with an average LFQi ratio of 1.5 across 3 biological replicates for AR and 2 biological replicates for SRF were chosen. The same method was applied to AR endogenous + DHT/siRNA and SRF vector + DHT/siRNA.

### STRING database search

A STRING (v11.5) database (<u>STRING: functional protein association networks</u> (string-db.org)) search was performed on MS hits to investigate protein-protein interaction. Using the ‘Multiple Proteins’ search bar, the hits associated with either SRF or AR were inputted into the search list (with additional input of AR and SRF also) and searched against the ‘*Homo sapiens’* database.

### 3-(4,5)-dimethylthiazol-2-yl-2,5-diphenlytetrazolium Bromide (MTT) cell viability assay

Three thousand cells per well were cultured in 96 well plates. Cells were treated with increasing concentrations of either JG-98, Ver-155008, Ganetispib, Ipatasertib or Alpelisib or DMSO control. Cells were treated for 5 days followed by MTT analysis as previously described^14^.

### Statistical Analysis

IC_50_ values were calculated using a non-linear regression dose response curve on Graphpad Prism version 7. Unpaired two-tailed Student’s t-tests were performed to compare treated conditions with the vehicle control. P values below 0.05 were considered statistically significant. All tests are indicated in the table legends.

## Results

### Interactome analysis identifies AR and SRF common interactors

The experimental design for this discovery approach is outlined in Figure 1. LNCaP Abl cells were transfected with scrambled siRNA, AR siRNA or SRF siRNA, and SRF vector in the presence or absence of 10nM DHT (for 24 hours) before pulldown. DHT stimulation was introduced to enrich for co-factors involved in transcriptional regulation of AR. The siRNA samples were used as a negative control to eliminate any non-specific interactors during data analysis. Overexpression of SRF enhanced binding of interactors that would be otherwise missed, while AR is already overexpressed in LNCaP Abl cells^11^. SRF and AR downregulation and SRF overexpression were confirmed by WB (Figure 2). 168 proteins for the AR pull down and 157 for the SRF pull down were identified before and after DHT combined (Supplementary tables 1-2). The data obtained from MaxQuant was analysed on Perseus software to remove contaminants and reverse proteins, which are generally defined as CrapOME, nonspecific for proteomic analysis^15^ and could be false positives or nonbiological protein sequences respectively. Proteins that did not appear in at least two of the three biological replicates were discarded. LFQi ratios of either AR endogenous (with and without DHT stimulation) *vs.* AR siRNA, or SRF overexpression *vs*. SRF siRNA (with and without DHT stimulation) were calculated for each independent experiment.

**Figure 1.**
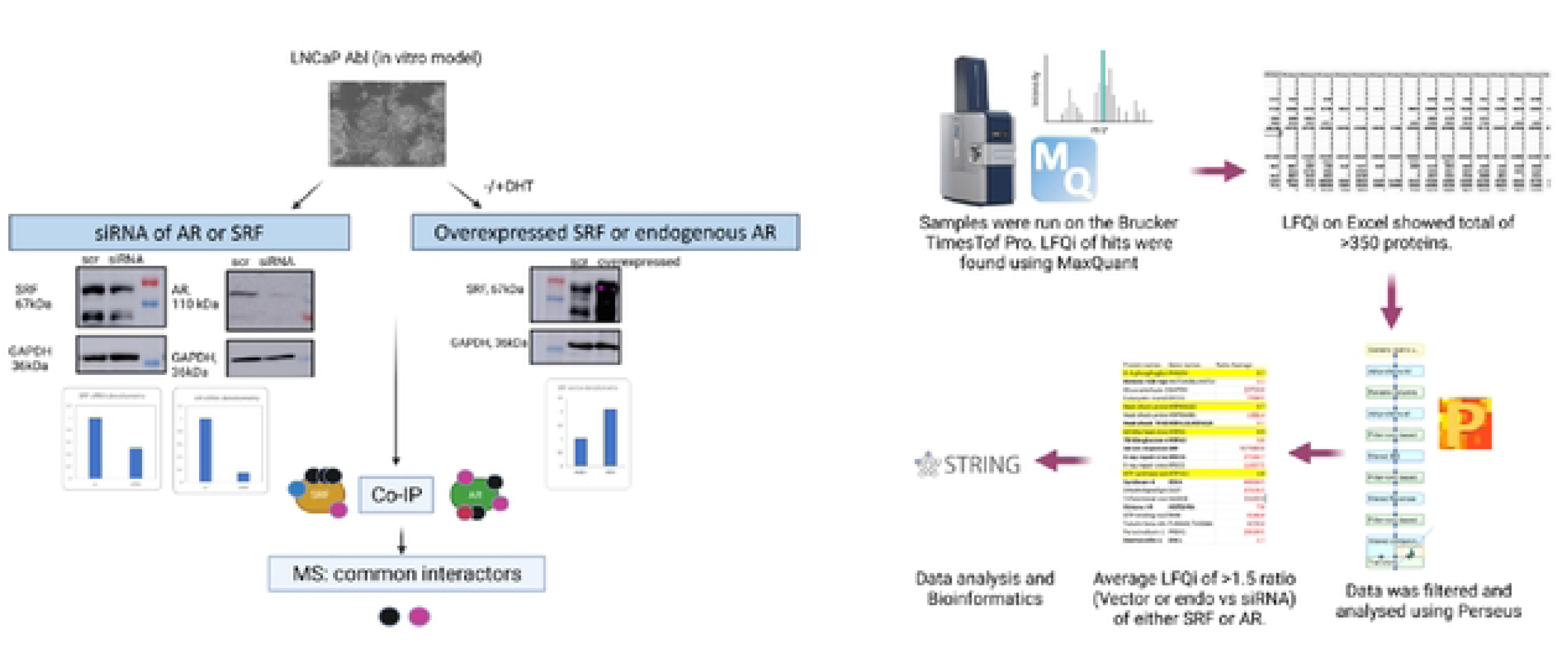
Discovery approach to identify the common interactors in the SRF and AR pathways. A. Schematic diagram of the experimental design. LNCaP Abl cells were transfected with either scramble siRNA, AR siRNA, SRF siRNA or with a plasmid overexpressing SRF, with or without DHT stimulation. LC-MS was performed following Co-IP pulldowns with antibodies specific for AR or SRF. B. Schematic diagram of data analysis following MS.

**Figure 2.**
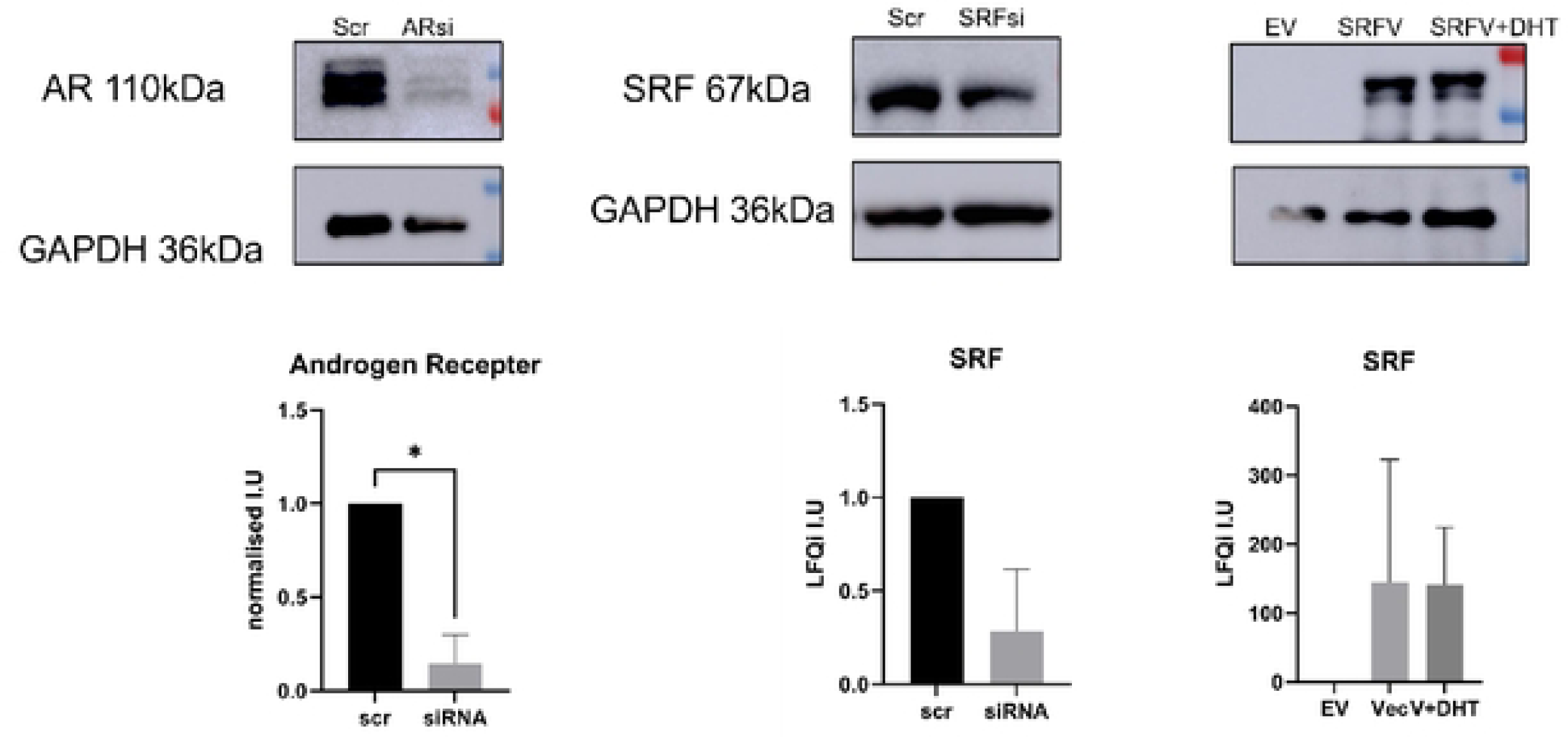
WB validation of downregulation of AR and SRF and upregulation of SRF following siRNA/plasmid transfections. Top panels show WB representative images of three independent experiments. Bottom panels show densitometry analysis of WB images. Bars represent average of three independent experiments ± standard deviation. Abbrev: Scr = scrambled, EV = empty vector, SRFV = SRF overexpressing vector, SRFV+DHT = SRF overexpressing vector + DHT. Two-tailed t-test was carried out followed by Welch’s correction. * = p<0.05

To identify the proteins with increased expression in the overexpressed (SRF) or endogenous (AR) sample compared to siRNA, LFQi ratios were used to obtain a cut-off of ≥1.5 of overexpression/siRNA or endogenous/siRNA, in line with previous proteomic studies on the AR interactome^16^. Twenty-one proteins interacting with SRF were identified before DHT and 14 post-DHT stimulation (Figure 3, panel A). From the AR Co-IP, 10 proteins were identified before DHT and 6 after DHT stimulation(Figure 3, panel B). AR and SRF datasets were then analysed for common interactors. A total of 7 proteins were identified in all datasets (Table 1). The 7 common interactors include Heat-shock 70kDa protein (HSP70), Heat-shock protein 90α (HSP90α) and Heat-shock protein 90β (HSP90β), 78 kDa glucose-regulated protein (HSPA5) and Heat-shock cognate 71 kDa (HSPA8), Glyceraldehyde 3-phosphatase (GAPDH) and the antioxidant enzyme peroxiredoxin-1 (PRDX1). Interestingly, 5 of these proteins interacted with both AR and SRF without DHT stimulation (HSP70, HSP0AA1, HSP90AB1, HSPA5, PRDX1), there were no common proteins associated with AR and SRF after DHT stimulation, and 2 proteins were in common to SRF and AR at different DHT stimulation status (GAPDH interacted with SRF -DHT and AR +DHT and HSPA8 interacted with SRF +DHT and AR -DHT).

**Figure 3.**
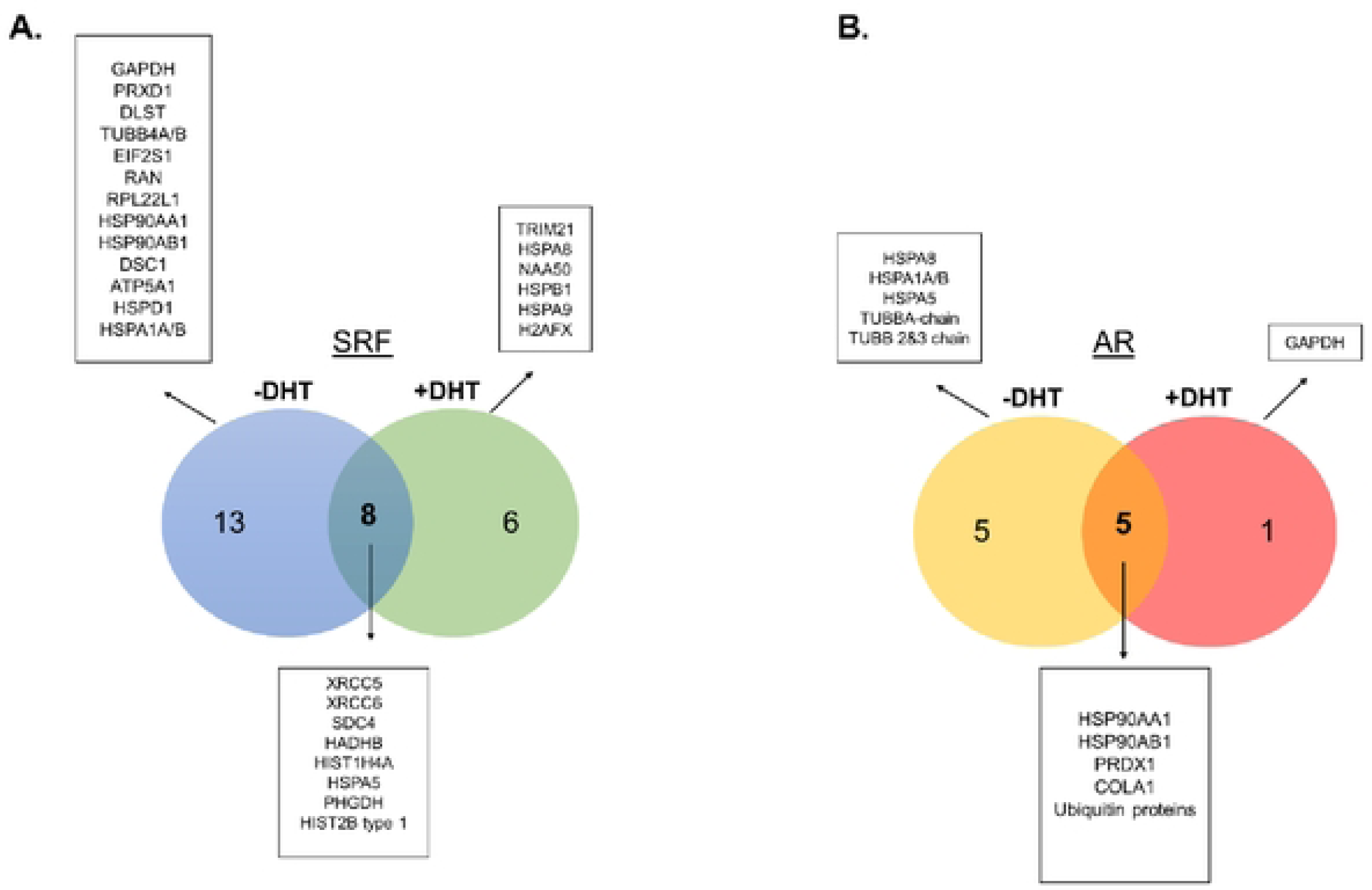
Summary of the identified proteins in different experimental groups. A) Proteins from the SRF dataset before and after DHT. B) Proteins from the AR dataset before and after DHT.

**Table 1.** List of AR/SRF common interactors, and their functions taken from Uniprot (Uniprot KB/Swiss-Prof).

| AR/SRF Common interactors |  |  |
| --- | --- | --- |
| Protein names | Gene names | Function (Uniprot Swiss-Prot) |
| Heat shock 70 kDa protein | HSPA1B;HSPA1A | Molecular Chaperone Protein involved in protein degradation and re-folding |
| Heat shock protein HSP 90-alpha* | HSP90AA1 | Chaperone protein that regulates proteins involved in cell cycle control and signal transduction |
| Heat shock protein HSP 90-beta | HSP90AB1 | HSP90 subunit |
| 78 kDa glucose-regulated protein | HSPA5 | Endoplasmic reticulum chaperone protein |
| Heat shock cognate 71 kDa protein* | HSPA8 | Molecular chaperone protein |
| Glyceraldehyde-3-phosphate dehydrogenase | GAPDH | Key enzyme in glycolysis |
| Peroxiredoxin-1 | PRDX1 | Protects cells from oxidative stress |

### Pathway analysis of the AR/SRF common interactors

The common interactors of AR/SRF were analysed using STRING to identify other pathways in the interactome that may be vulnerable to inhibitors (Figure 4, panel A). This analysis identified proline, glutamate and leucine rich protein 1 (PELP-1), a steroid nuclear receptor adaptor protein that has known interactions with both AR and SRF^17^. Additionally, CTNNB1 (or β-catenin), a major component of the WNT signalling pathway, has been shown to interact with AR and weakly associated with SRF^17,18^. Using the expansion method on the STRING database with the common AR/SRF interactors as the input, enriched pathways were investigated, using the KEGG database. PI3k-Akt and MAPK signalling pathways resulted significantly enriched in this analysis(Figure 4, panel B).

**Figure 4.**
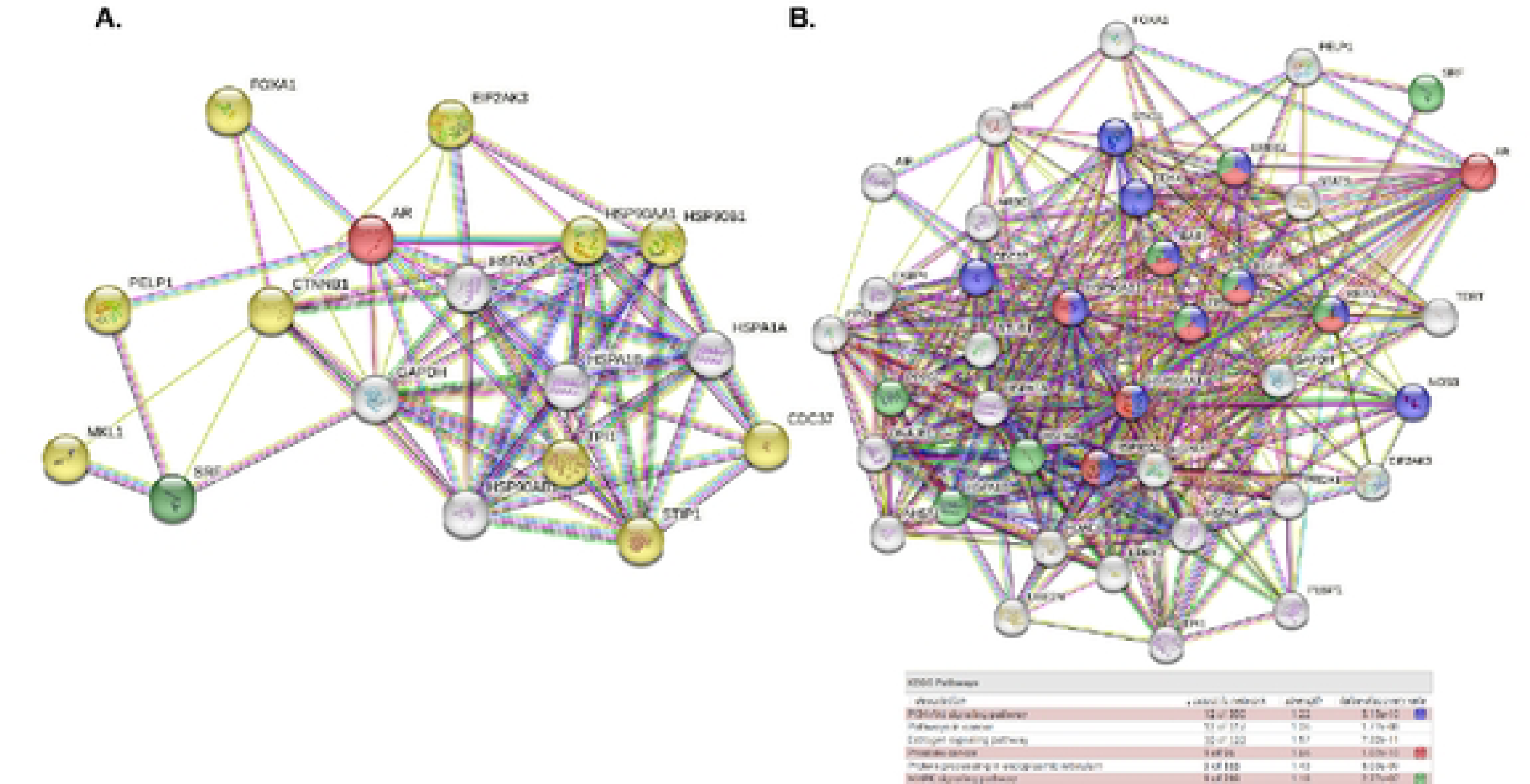
Pathway and network analysis of the proteins identified in the ARISRF interactome. STRING analysis of the common interactors between SRF and AR revealed other proteins involved in the ARISRF interactome. A. Common interactors and their known interactions with AR and SRF. Common interactor nodes are shown in white. AR and SRF nodes are shown in red and green respectively. Yellow nodes represent other proteins following network expansion. BJ Following network expansion, a list of highly significant KEGG pathways were highlighted. Blue nodes represent proteins involved in the Pl3k-Akt pathways, red nodes proteins involved in prostate cancer signalling, and green nodes MAPK signalling pathway. The highlighted pathways all have a false discovery ratio below 0.05.

### Functional validation of proteins identified in the AR/SRF interactome

The MS data coupled with bioinformatic analysis identified several interesting targets, summarised in Table 2. HSP70, HSP90 and AKT signalling were prioritised for further functional validation. The drugs tested included the HSP70 inhibitors, Ver-155008 and JG-98, HSP90 inhibitor Ganetespib, PI3k inhibitor Alpelisib and AKT inhibitor Ipatasertib. The rationale for starting our functional investigation from these three targets is as follows. Inhibition of HSP70 in pre-clinical models of CRPC has shown promise in its capability as therapeutic target and in overcoming resistance to AR antagonists^20^. HSP90 is a known AR coregulator^21^ that is regulated by HSP27, which contains Serum Responsive Elements in its promoter region, and therefore is potentially under the transcriptional control of SRF^22^, making it an ideal target in the context of our study. Moreover, HSP90 inhibition in combination with ADT is currently being investigated in clinical trials (Trials: NCT0168526 and NCT01270880). One of these trials include the HSP90 inhibitor, Ganetespib, which was tested in patients with CRPC^23^ but was not deemed efficient as a single drug, therefore our study may offer novel avenues for combination treatments. As the PI3k/AKT signalling is upstream of both AR and SRF and emerged as an enriched pathway in the STRING analysis, we also tested Ipatasertib and Alpelisib. Ipatasertib is an AKT inhibitor that is in phase III clinical trials in combination with abiraterone acetate for mCRPC (NCT03072238) and for CRPC patients that developed resistance to docetaxel (NCT01485861)^24^. Among the numerous PI3k inhibitors currently available, we selected Alpelisib, which targets the *PI3KCA* mutation, associated with poor survival in patients with PCa^25^.

**Table 2.** List of molecular targets identified through MS and STRING analysis.

| Target | Gene ID | Function | Inhibitor(s) | Clinical Trial ID |
| --- | --- | --- | --- | --- |
| Heat Shock protein 90 | HSP90AA1<br>HSP90AB1 | AR co-regulator and chaperone protein | Onalespib<br>Ganetespib<br>Luminespib | NCT01685268 (met PCa with ADT)<br>NCT01270880 (met PCa) |
| HSP70 family of proteins | HSPA1/B;<br>HSPA5; HSPA9 | Chaperone proteins | VER-155008<br>JG-98 series of HSP70 inhibitors | Pre-clinical models |
| Ku70/80 | XRCC5 and<br>XRCC6 | AR co-regulator, DNA repair protein | Vitas-M STL127705 | n/a |
| Tubulin beta chain IV and chain II | TUBB4 and<br>TUBB2 | Constituent of microtubules | Tubulin Inhibitors (eg docetaxel, cabazitaxel, vinblastine, | Many clinical trials with docetaxel and cabazitaxel for met PCa.<br>ABT-751 |
| Peroxiredoxin-1 | PRDX1 | Antioxidant enzyme | PRDX1 Inhibitor H7 | n/a |
| PELP-1 | PELP1 | AR co-regulator, adaptor protein | Peptidomimetics* | Preclinical models |
| PI3k complex | PIK3CA | Activates multiple intracellular signaling pathways | Alpesilib | n/a |
| Akt | AKT1 | Tyrosine kinase pathway | Ipatasertib<br>ONC-201 (ERK/Akt inhibitor) | NCT03072238 (met PCa with ADT)<br>NCT01485861 (met PCa)<br>Phase 3 GBM |

Following 5 days of treatment viability was assessed using MTT assays. In the Parental cells, IC_50_ concentrations were as follows: 10.3μM±1.7 for Ver-155008, 0.13µM±SD for JG-98 and 17.2nM±7.9nM for Ipatasertib (Figure 5, panel A, Table 3). Despite both drugs target HSP70, the IC_50_ concentration of JG-98 was lower than that of Ver-155008, possibly due to different mechanisms of action of these drugs^24^. In the LNCaP Abl cells, the IC_50_ concentrations of Ver-155008 was 12.5µM±4.1, JG-98 was 70nM±30, Ipatasertib was 25.6nM±13.6 and Ganetespib was 18.7nM±6.7 (Figure 5, panel B, Table 4). Interestingly, the IC_50_ values of JG-98 and Ipatasertib were significantly lower than that of enzalutamide previously reported in LNCaP parental cells (8.8µM±3.4)^14^. In the Abl cell line, which models CRPC, the IC_50_ values of Ver-155008, JG-98, Ipatasertib and Ganetespib were also significantly lower than that of enzalutamide (26.3µM±6.9).

**Figure 5.**
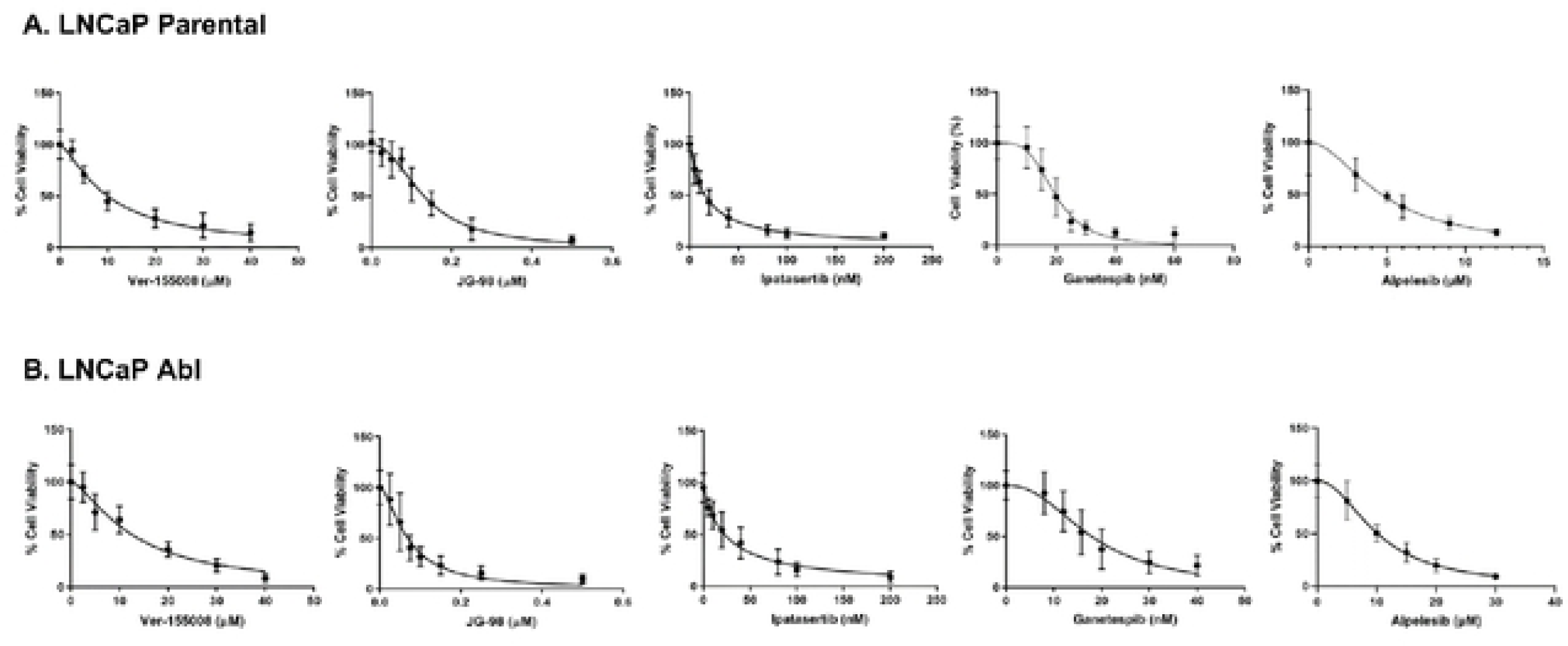
Cell Viability graphs of inhibitors against HSPTO, HSP90 P/3k and AKT in LNCaP parental and Abl cells. Dose dependent curves of inhibitors Ver-155008, JG-98, lpatasertib, Ganetespib and Alpelesib. A. LNCaP Parental. Concentrations used are as follows: Ver-155008 (µM): 0, 2.5, 5, 10, 20, 30, 40. JG-98 (µM): 0, 0.025, 0.050, 0.075, 0.1, 0.15, 0.25, 0.5. lpatasertib (nM): 0, 5, 10, 20, 20, 40, 80, 100, 200. Ganetesib (nM): 0, 10, 20, 15, 20, 30, 40, 60. Alpelisib (µM): 0, 3, 6, 9, 12 B) LNCaP Abl. Concentrations used are as follows: Ver-155008 (µM): 0, 2.5, 5, 10, 20, 30, 40. JG-98 (µM): 0, 0.025, 0.05, 0.075, 0.1, 0.15, 0.25, 0.5. lpatasertib (nM): 0, 5, 10, 20, 40, 80, 100, 200. Ganestespib (nM): 0, 8, 12, 15.8, 20, 30, 40. Alpelesib (µM): 0, 1, 5, 15, 20. For each inhibitor, graphs represent the average of at least three biological replicates in triplicate. Error bars represent standard devation.

**Table 3.**
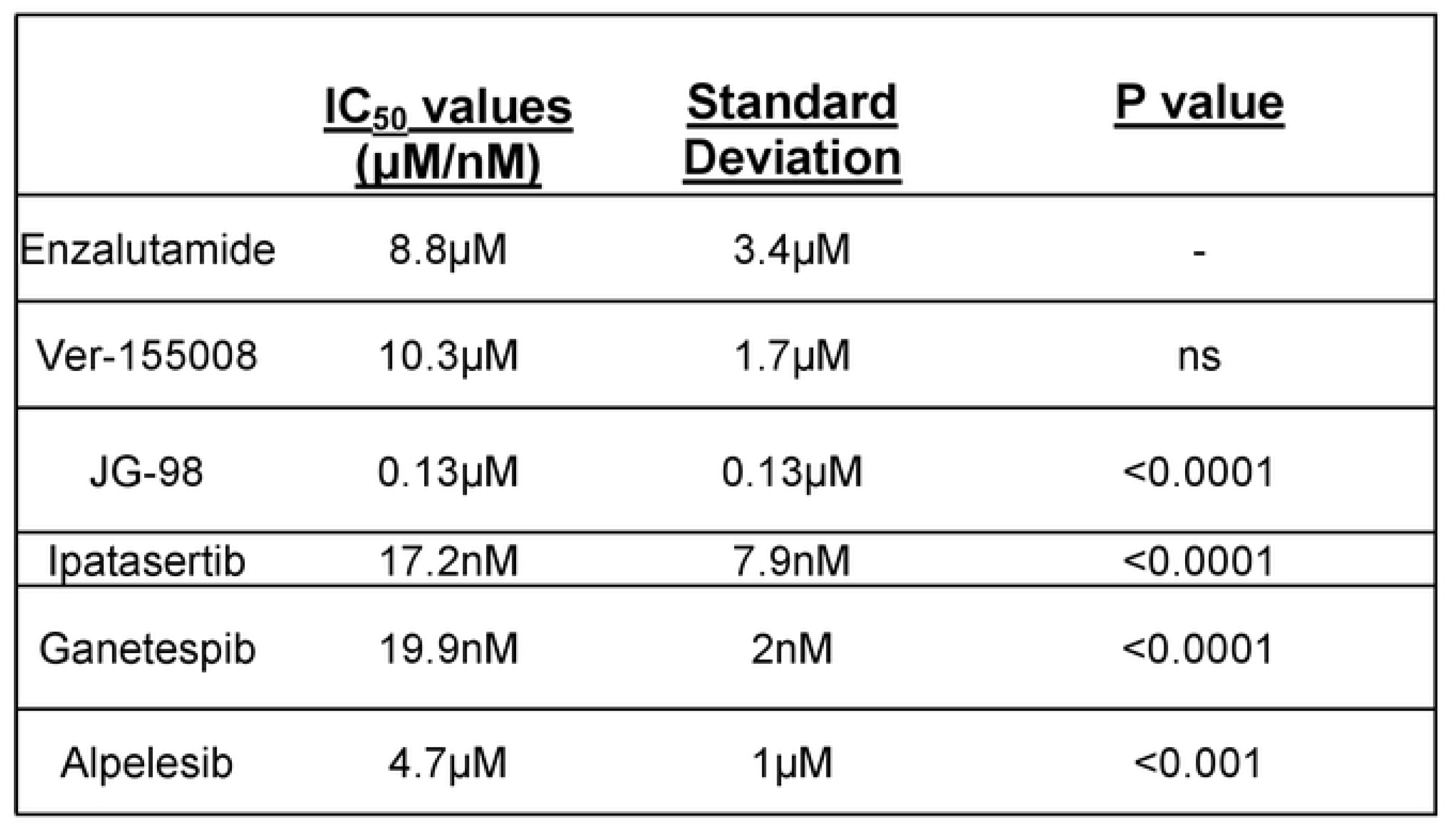
IC_50_ values of inhibitors compared against enzalutamide in LNCaP Parental cells.

**Table 4.** IC_50_ values of inhibitors compared against enzalutamide in the LNCaP Abl cells.

|  | <u>IC<sub>50</sub> values</u><br><u>(μM/nM)</u> | <u>Standard</u><br><u>Deviation</u> | <u>P value</u> |
| --- | --- | --- | --- |
| Enzalutamide | 26.3μM | 6.9μM | - |
| Ver-155008 | 12.5μM | 4.1μM | <0.0001 |
| JG-98 | 70nM | 30nM | <0.0001 |
| Ipatasertib | 25.6nM | 13.6nM | <0.0001 |
| Ganetespib | 18.7nM | 6.7nM | <0.0001 |
| Alpelesib | 19.18μM | 1μM | <0.0001 |

## Discussion

In this study 21 proteins were detected in the SRF Co-IP before DHT stimulation and 14 after. Among them there were XRCC5 and XRCC6, also known as Ku80 and Ku70, heterodimeric DNA repair proteins involved in non-homologous end joining of dsDNA breaks in the cell^26^. Ku70/80 are known AR co-activators and aid in its nuclear translocation in LNCaP cells, particularly in the presence of DHT. Their interaction with SRF, a master regulator of actin filaments^22^, happens through polymerised F-actin, which is vital for Ku70 localisation and Ku80 stabilisation during dsDNA breaks^26^. Another AR known coactivator that co-precipitated with SRF is RAN, also known as ARA24. RAN is a nucleocytoplasmic protein that directly binds to the NH2-COOH transcription activating domain of the AR protein^27^. Significantly increased RAN expression in PCa samples compared to normal adjacent samples was demonstrated^51^. Additionally, RAN colocalization with AR was predominantly observed in the nucleus and was enhanced in the presence of DHT, suggesting that RAN is particularly important in AR nuclear compartmentalisation and transcriptional activation^27^.

Pathway analysis of the MS hits of the AR/SRF interactome, lead to other potential common interactors, including the AR coactivator and adaptor protein PELP-1 and β-catenin. PELP-1 interaction with SRF was previously shown in NIH3T3 cells, where PELP-1 acts as a corepressor of SRF transcriptional activity^28^. PELP-1 is also a coregulator of nuclear steroid transcription factors and is involved in cell cycle progression, playing an oncogenic role in breast and prostate cancer^29^. Supporting our STRING analysis, a study of the AR interactome showed that PELP-1 and β-catenin were significantly upregulated in PCa tumour compared to normal tissue^30^. Despite its large molecular weight, peptidomimetics against PELP-1 in primary PCa explants and PCa xenografts have shown promise in disrupting AR translocation to the nucleus, resulting in decreased AR-mediated transcription^28^.

The PI3k/AKT pathway emerged as the most enriched pathway following expansion of STRING analysis when all the 7 common interactors were inputted. This pathway is aberrant in many different cancers and leads to enhanced cell proliferation and resistance to apoptotic cell death^24^. Inhibitors of AKT such as Capavisertib and Ipatasertib have shown promise as combination treatment with ADT in patients with CRPC and PTEN loss^24^. Importantly, a crosstalk has been established between AR and the PI3k/AKT pathway in PCa, with the PI3k/AKT pathway activated in 42% of localised PCa cases and 100% in advanced forms of disease^24^. Furthermore, inhibition of AR leads to overexpression of the PI3k/AKT pathway leading to aggressive growth, metastases and resistance to treatments. A downstream target of this pathway, mTOR, was shown to regulate the expression of HMMR in PCa cells via SRF, suggesting a possible link between the PI3k/AKT/AR/SRF pathways in PCa. Overall, we propose four possible common pathways in the SRF/AR interactome as detailed in Figure 6.

**Figure 6.**
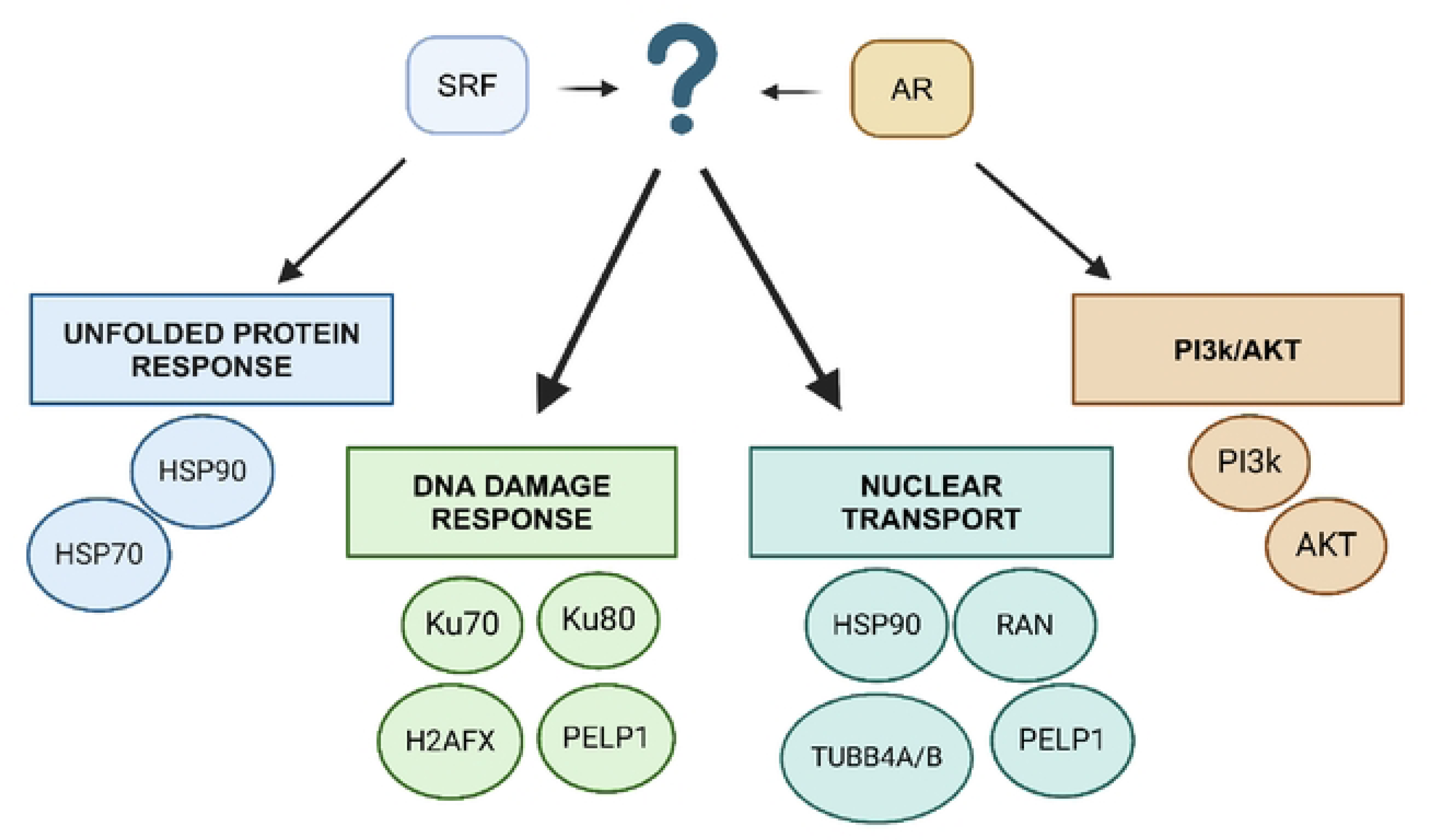
Proposed common pathways of the SRF/AR interactome and the key proteins identified by APMS and bioinformatics.

Out of our list of possible targets in the AR/SRF pathway, HSP90, HSP70, AKT and PI3K were functionally validated using Ganetespib, VER-155008/JG-98, Ipatasertib and Alpelesib respectively. Our results verified that inhibition of these proteins reduced cell viability in both LNCaP parental and Abl cells. Their IC_50_ values were significantly lower than that of enzalutamide, suggesting promise for further investigation. Specifically, combinations of inhibitors of key proteins in the AR/SRF signalling pathways with current therapies targeting AR and SRF inhibitors, may result in synergy in decreasing cell viability and proliferation, limiting PCa progression.

## Supplemental Materials

**Supplemental table 1.**
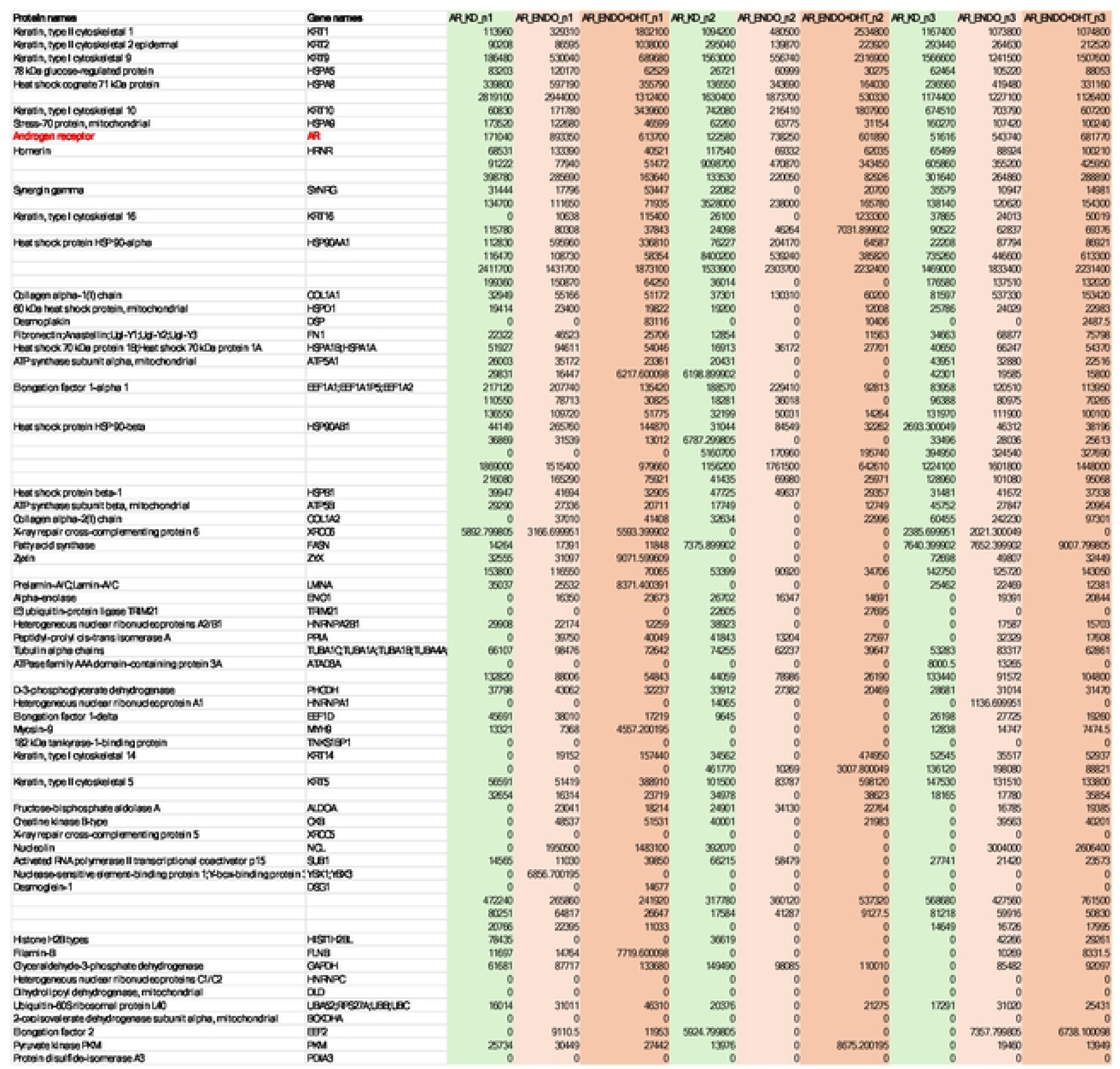
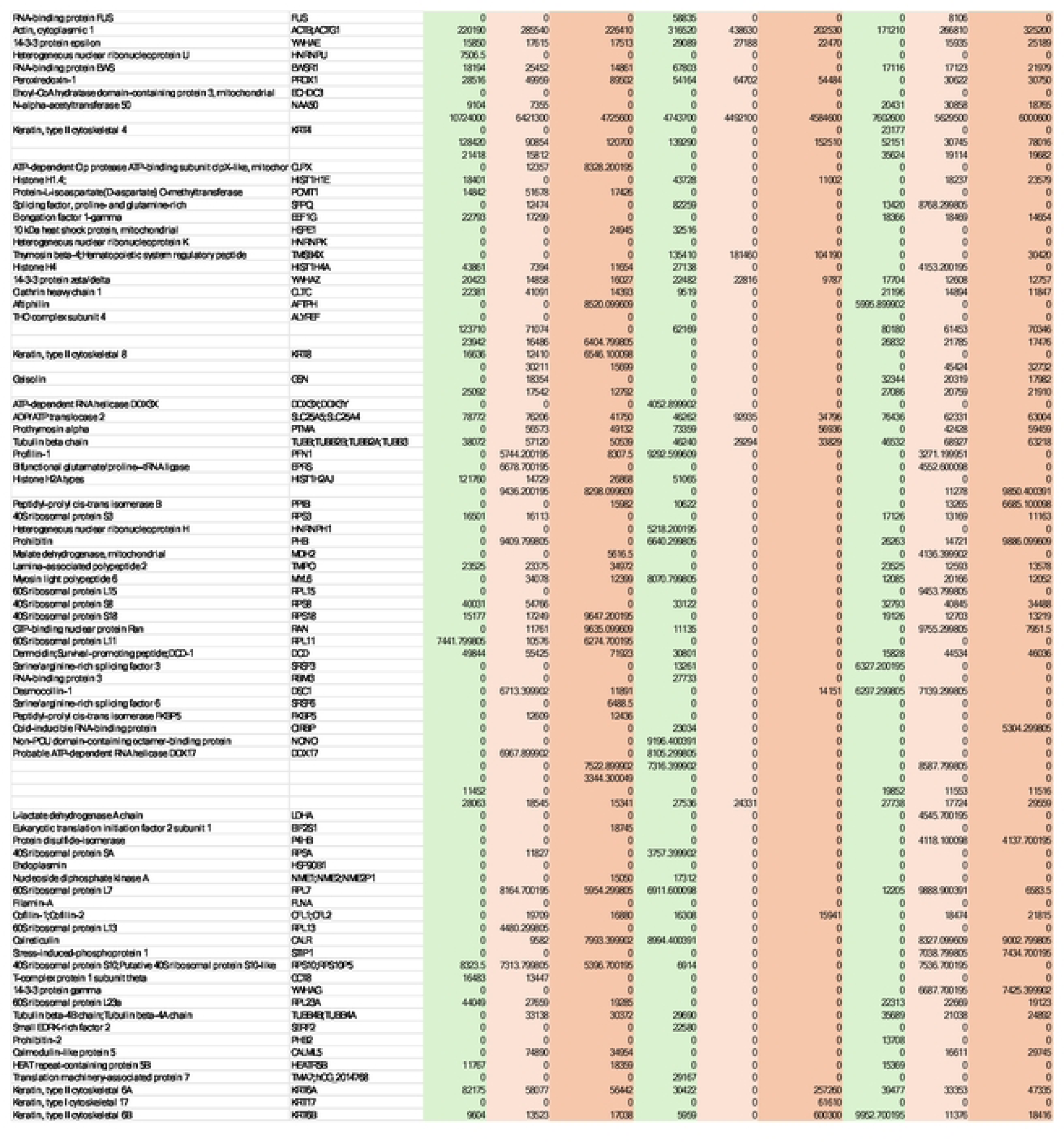
List of peptides precipitated with AR. Numbers indicate LFQi values. Abbreviations: AR KD, AR knocked down; AR ENDO, endogenous AR;

**Supplemental table 2.**
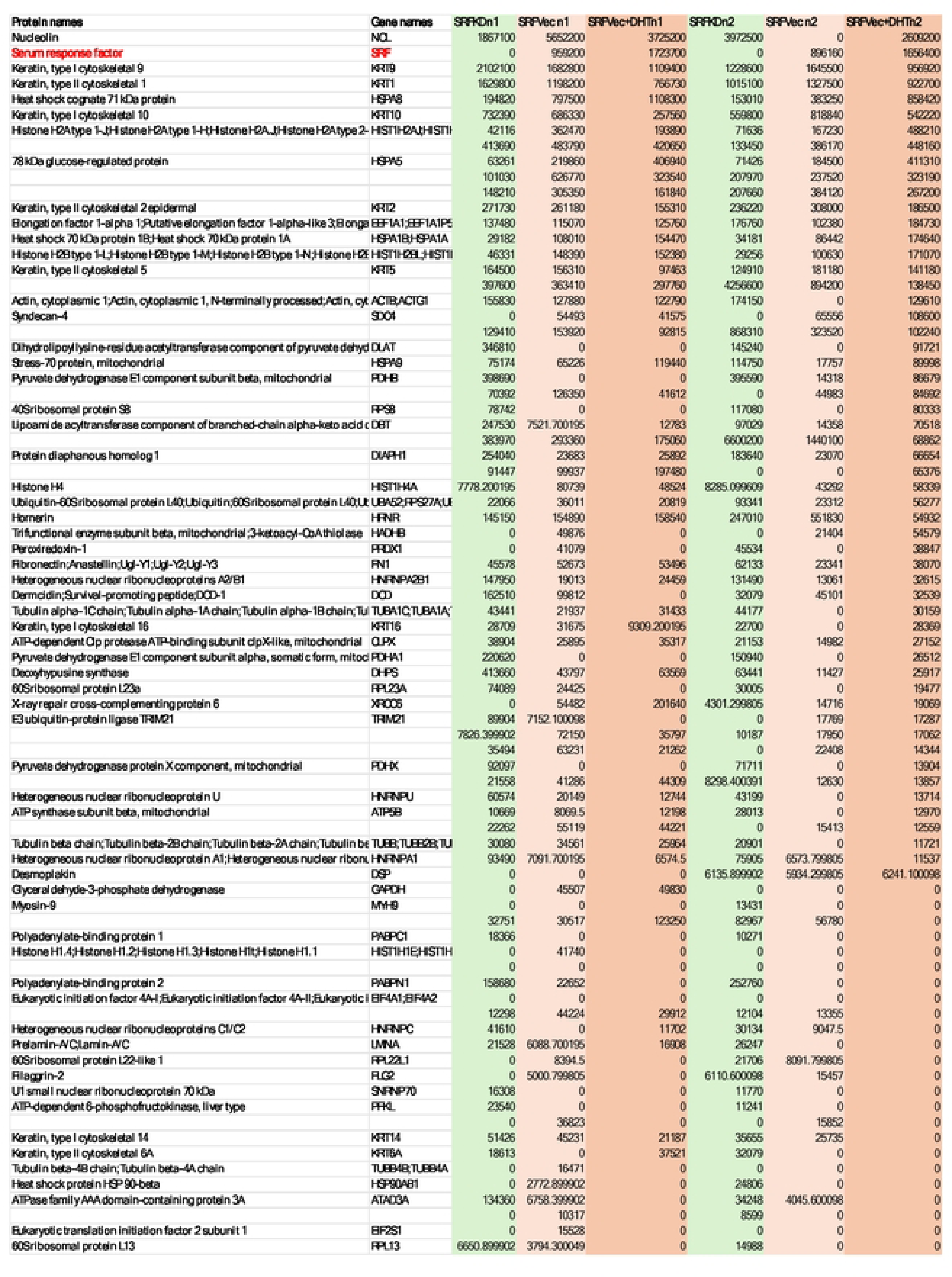
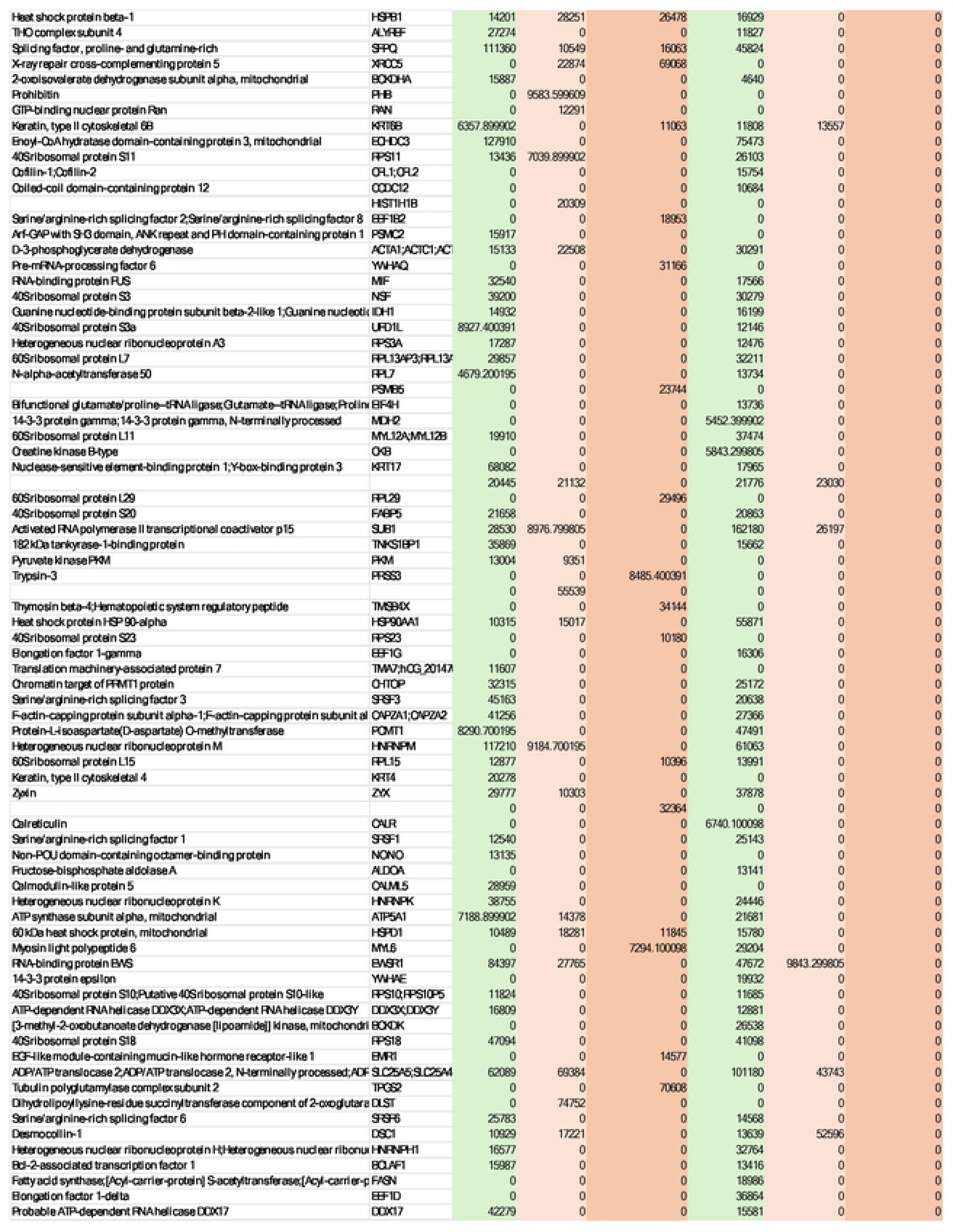
List of peptides precipitated with SRF. Numbers indicate LFQi values. Abbreviations: SRF KD, SRF knockdown; SRF Vec, SRF upregulation vector.

